# The essential chaperone DNAJC17 activates HSP70 to coordinate RNA splicing and G2-M progression

**DOI:** 10.1101/2023.10.25.564066

**Authors:** David V. Allegakoen, Kristen Kwong, Jacqueline Morales, Trever G. Bivona, Amit J. Sabnis

**Affiliations:** Department of Pediatrics, Division of Oncology, University of California San Francisco, San Francisco CA; Department of Medicine, Division of Hematology-Oncology, University of California San Francisco, San Francisco CA; Chan Zuckerberg Biohub, San Francisco CA

## Abstract

Molecular chaperones including the heat-shock protein 70-kilodalton (HSP70) family and the J-domain containing protein (JDP) co-chaperones maintain homeostatic balance in eukaryotic cells through regulation of the proteome. The expansive JDP family helps direct specific HSP70 functions, and yet loss of single JDP-encoding genes is widely tolerated by mammalian cells, suggesting a high degree of redundancy. By contrast, essential JDPs might carry out HSP70-independent functions or fill cell-context dependent, highly specialized roles within the proteostasis network.

Using a genetic screen of JDPs in human cancer cell lines, we found the RNA recognition motif (RRM) containing *DNAJC17* to be pan-essential and investigated the contribution of its structural domains to biochemical and cellular function. We found that the RRM exerts an auto-inhibitory effect on the ability of DNAJC17 to allosterically activate ATP hydrolysis by HSP70. The J-domain, but neither the RRM nor a distal C-terminal alpha helix are required to rescue cell viability after loss of endogenous *DNAJC17*. Knockdown of *DNAJC17* leads to relatively few conserved changes in the abundance of individual mRNAs, but instead deranges gene expression through exon skipping, primarily of genes involved in cell cycle progression. Concordant with cell viability experiments, the C-terminal portions of *DNAJC17* are dispensable for restoring splicing and G2-M progression.

Overall, our findings identify essential cellular JDPs and suggest that diversification in JDP structure extends the HSP70-JDP system to control divergent processes such as RNA splicing. Future investigations into the structural basis for auto-inhibition of the DNAJC17 J-domain and the molecular regulation of splicing by these components may provide insights on how conserved biochemical mechanisms can be programmed to fill unique, non-redundant cellular roles and broaden the scope of the proteostasis network.

## Introduction

Heat shock proteins are a diverse class of molecular chaperones that support the synthesis, folding, trafficking, and regulated destruction of proteins. Heat shock protein-70 kiloDalton (HSP70) proteins are an evolutionarily conserved family of chaperones that utilize ATP hydrolysis to maintain the protein homeostatic networks of both prokaryotic and eukaryotic cells. A class of co-chaperones, the heat shock protein-40 kiloDaltons (HSP40) or J-domain proteins (JDP), have evolved alongside the HSP70 family in metazoans to enable diverse, tightly regulated cellular functions, with the basic mechanics of their allosteric interactions largely conserved from bacterial DnaJ and DnaK, respectively.^1^ Central to this interaction is the J-domain, which permits acceleration of ATP hydrolysis by DnaK/HSP70, and is the defining feature of JDP members. Nonetheless, some JDP exhibit essential J-domain independent functions.^2^ This may point either to HSP70-independent cellular activities, or result from redundancy that permits other related JDPs to compensate for loss of any single unit.^3^

While the housekeeping roles of the collective ‘chaperome’ are broadly essential for cellular survival, the unique demands of different cell states can create cell type-selective chaperone dependencies. As a proof of concept, we demonstrated that genomically diverse subsets of rhabdomyosarcoma (RMS), the most common soft tissue cancer of childhood, harbor a selective dependence on cytosolic HSP70 to suppress ER stress responses and enable cancer progression.^4^ A corollary hypothesis arising from this finding is that individual JDPs might similarly carry out functions that are essential in some contexts, but not others. Alternatively, the functions of specific JDP-HSP70 pairs may be broadly essential due to a specialization in function that cannot be compensated for by the proteostasis network.

Here, we harnessed genetic screens to test the necessity of different JDPs in pediatric cancer cell lines, and carried out structure-function studies of the essential HSP70 co-chaperone DNAJC17. Through *in vitro* and cellular analysis, we find that the domains required for DNAJC17 to support cell survival connect HSP70 activation with mRNA splicing, and uncover dispensable regulatory domains that fine tune this protein’s activity.

## Results

### *DNAJC17* is an essential JDP in human cancer cell lines

We sought to test the hypothesis that selective HSP70 dependence in rhabdomyosarcoma is due to the cellular effects of a unique HSP70-JDP pair. To do so, we conducted a CRISPRi^5^ screen of JDP family members in the patient derived Rh30 cell line. Cells expressing the dCas9-KRAB transcriptional repressor were infected with a pooled lentiviral library, and deep sequencing was used to quantify sgRNA abundance immediately following puromycin selection (t=0) and after ten days in cell culture (t=10). Across 41 different JDP, we found that guides targeting only two genes, *DNAJC17* and *DNAJA3*, had strong and consistent depletion (**Figure 1A**).

**Figure 1.**
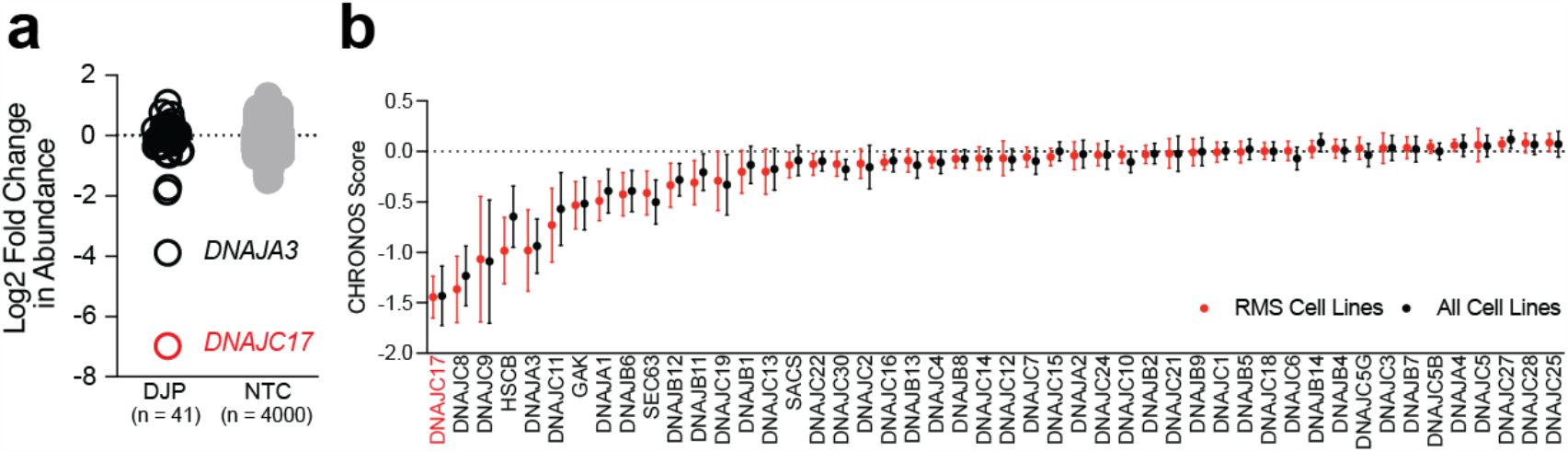
DNAC17 is a pan-essential member of the JDP family. A, Depletion of sgRNA targeting 41 distinct JDPs or 4000 non-targeting controls (NTC) over 10 days in the Rh30 cell line from a pooled CRISPRi screen. Genes with depletion over NTCs include *DNAJC17* and *DNAJA3*. B, data from the DepMap project^6^ showing mean and standard deviation of CHRONOS estimates of fitness effects for knockout of each of the indicated JDP-encoding genes. Scores from the 13 rhabdomyosarcoma cell lines are shown in red, compared to scores from all 1,078 cell lines in black.

We next leveraged the Dependency Map (DepMap) project^6^ to ask whether dependencies for *DNAJC17* or *DNAJA3* were unique to rhabdomyosarcoma. Unexpectedly, *DNAJC17* stood out as the most essential JDP not just in RMS, but across all cell lines in the DepMap project (although no unique guide RNAs are included in the DepMap dataset to interrogate *DNAJB3*) (**Figure 1B**). Confirming the functional redundancy of the proteostasis network, only 3 out of 48 JDP-encoding genes in the DepMap had a CHRONOS score of less than -1, indicating essentiality. We note that the DepMap can identify selective proteostasis dependencies; consistent with our previous work, only 167 out of 1078 cell lines had a CHRONOS dependency score of less than -1 for the HSC70-encoding gene *HSPA8*, and it is the only HSP70-encoding gene that shows evidence of RMS selectivity (p < 0.0001 by two-way ANOVA with post-hoc Sidak’s test; **Supplementary Figure S1**). A similar analysis showed no enhanced RMS cell dependence on the majority of JDP-encoding genes. *HSCB*, a co-chaperone involved in iron-sulfur cluster biosynthesis, did show enrichment in RMS cells in the DepMap, but is still annotated as pan-essential and favored to interact specifically with the pan-essential mitochondrial HSP70 mortalin (*HSPA9*).^7^ In the current work, we turned our attention to understanding the outlier essential role of *DNAJC17*.

JDPs have been historically classified based on structurally conserved elements. Members of the third class, to which *DNAJC17* belongs, harbor an HSP70-interacting DNAJ domain but have wide sequence divergence from one another aside from this feature.^8^ The predicted protein structure of DNAJC17 includes a J-domain at its N-terminal end and an annotated RNA-binding RRM motif at its C-terminal end. The intervening segments of the protein are annotated as intrinsically disordered by MobiDB-Lite,^9^ but harbor a putative nuclear localization sequence (NLS). Computational prediction of DNAJC17’s structure by AlphaFold^10^ suggests that the linker region between the J-domain and RRM consists of a coiled coil, and also identifies a helical tail distal to the RRM (**Figure 2A-B**).

**Figure 2.**
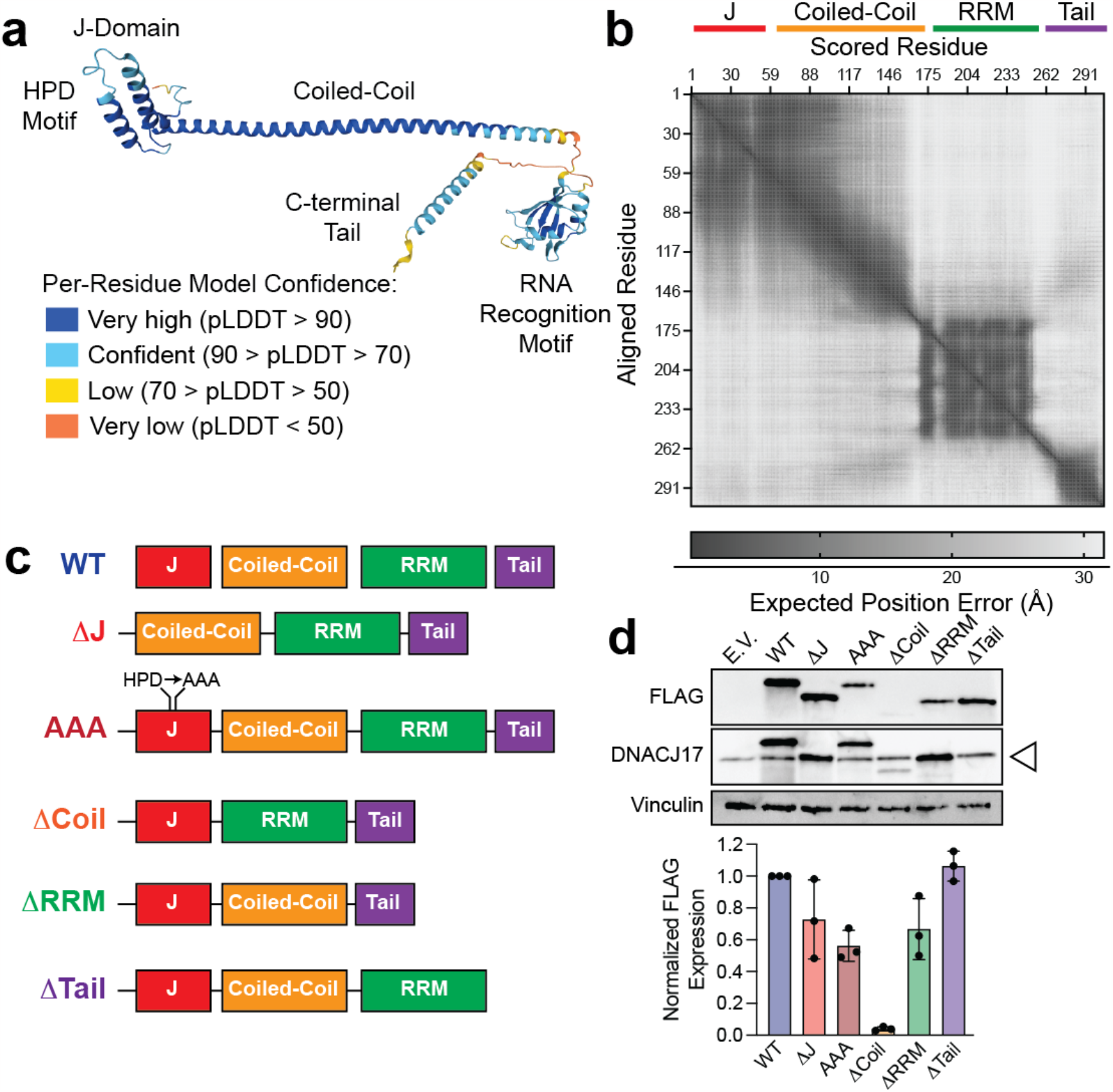
Deletion mutants of DNAJC17 to shed light on structure-function relationships. Prediction of DNAJC17 structure from AlphaFold shown as a structural model (A), or through expected position error of amino acid alignments relative to one other (B). C, schematic of different alleles used to study the relationship of DNAJC17 structural domains to its biochemical and cellular functions. D, representative immunoblot and quantification of the expression of FLAG-tagged constructs transiently transfected into HEK-293 cells. Open triangle shows endogenous DNAJC17, which as approximately the same molecular weight as the ΔJ and ΔRRM mutants. Barplots depict the mean, and error bars show standard deviation.

We sought to use these four structural elements (J-domain, coiled-coil, RRM, and C-terminal tail) as a starting point to better understand the essential roles of DNAJC17 in cancer cells (**Figure 2C**). We generated FLAG-tagged deletion alleles, as well as a missense mutant allele of the conserved HPD motif in the J-domain,^11^ to do so. We transiently transfected Rh30 cells with equal amounts of FLAG-tagged plasmids, and found that wildtype DNAJC17 levels consistently exceeded other mutants, with similar levels of the ΔJ, ΔRRM, and ΔTail constructs (**Figure 2D**). Conversely, the ΔCoil mutant had extremely low expression, and was not further analyzed based on its instability for cellular assays.

### DNAJC17 harbors an active J-domain that is inhibited by the RRM

Prior studies of an RNA-binding *DNAJC17* ortholog in yeast, *Cwc23*, found that the J-domain of that protein was dispensable for its function.^12^ As a first step to understand the necessity of the J-domain of *DNAJC17*, we sought to confirm that it is biochemically functional. The J-domain of DNAJC17 shows a high degree of homology with the J-domain of DNAJB1 and *E. coli* DnaJ (**Figure 3A**), including 47% and 39% identity and 78% and 67% positives. We purified codon-optimized wildtype DNAJC17 and tested its capacity to accelerate ATP hydrolysis of purified HSP72, measured by malachite green measurement of free phosphate release.

**Figure 3.**
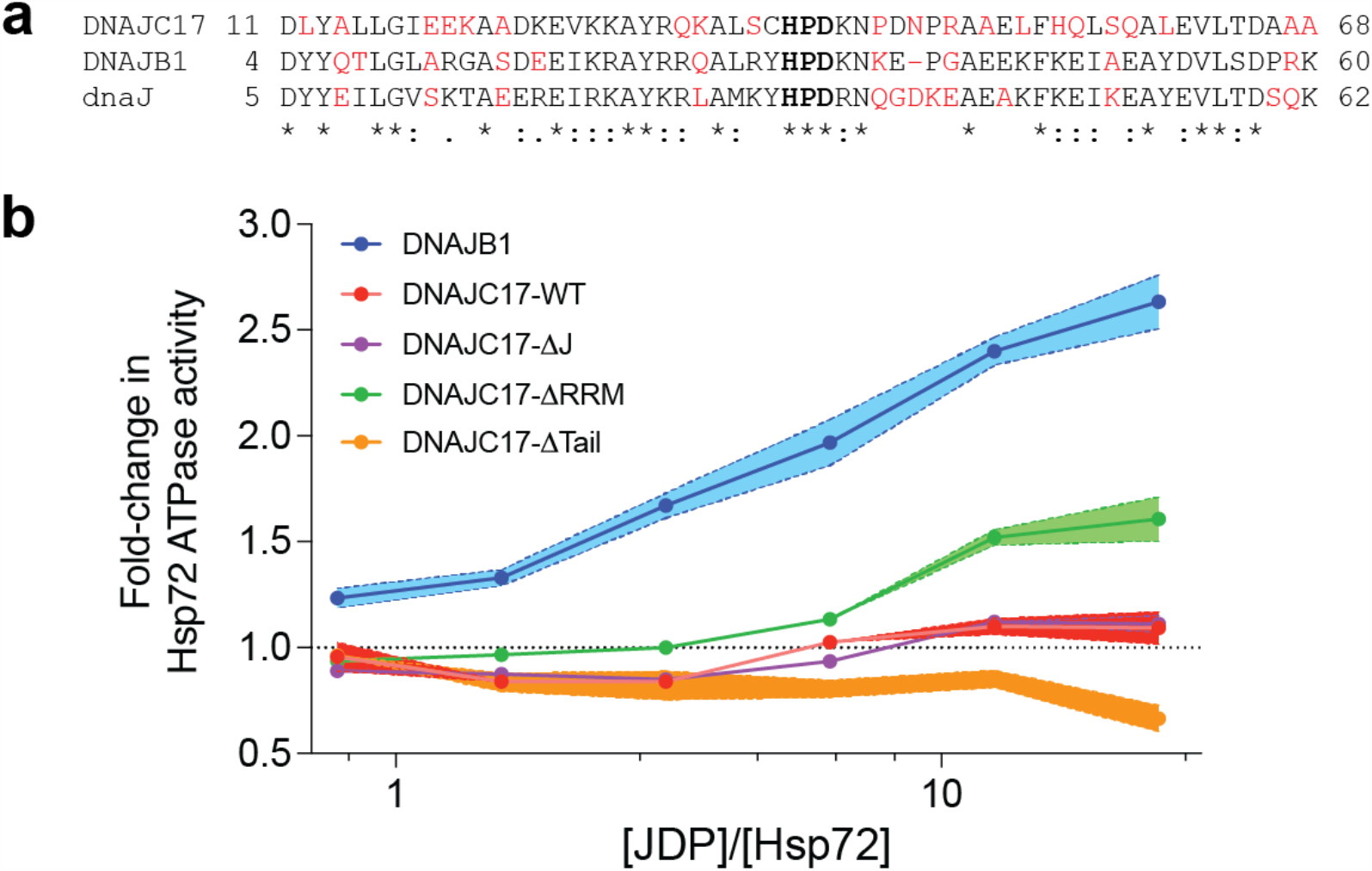
The J-domain of DNAJC17 can stimulate ATP hydrolysis by HSP70 after relief of auto-inhibition. A, sequence alignment of the J-domains of *H. sapiens* DNAJC17, DNAJB1, and *E. coli* dnaJ. B, fold-change in ATP hydrolysis by recombinant HSP70 when incubated with the indicated concentrations of DNAJB1 or DNAJC17 mutants over 1 hour. Shaded regions depict standard error of the mean from three independent replicates.

Surprisingly, we found that purified DNAJC17 was unable to stimulate ATP hydrolysis by HSP72 (**Figure 3B**): ATPase activity was identical between wildtype and ΔJ DNAJC17. In class 2 JDPs, C-terminal domains can exert auto-inhibitory effects on J-domain function.^13^ Thus, we asked whether deletion of C-terminal domains might restore the activity of DNAJC17. Indeed, deletion of the C-terminal RRM (though not the distal tail) was sufficient to enable acceleration of ATP hydrolysis by HSP72, though still quite modest compared to DNAJB1. We conclude that the J-domain of DNAJC17 can induce allosteric activation of HSP70, and that the RRM attenuates this productive interaction.

DNAJC17 is predicted to bind RNA through its RNA recognition motif (RRM).^14,15^ However, attempts to isolate RNAs bound to either endogenous or overexpressed FLAG-tagged DNAJC17 were unsuccessful. Cells were lysed, subjected to FLAG-immunoprecipitation, and then total RNA was extracted and quantified through capillary electrophoresis. Given the described proteomic association of DNAJC17 with nucleolar splicing machinery,^16^ we included the RNA-binding spliceosome component SNRPA1 as a positive control. While SNRPA1 showed clear enrichment for small RNAs, electrophoretograms of DNAJC17 pulldowns were similar to negative controls (**Supplementary Figure S2**). The RRM may thus enable transient RNA interactions, or potentially enable protein-protein interactions.^17^ The specific cellular interactors of the DNAJC17 RRM thus remain to be identified.

### The N-terminal J domain of DNAJC17 is essential for G2-M progression

Having established a clear role for the J-domain and inhibitory RRM in allosteric interaction with HSP70, we next moved to cellular systems to understand how the structural domains of DNAJC17 dictate its function in sarcoma cells. We investigated the effects of *DNAJC17* loss in Rh30 rhabdomyosarcoma cells, and as a parallel, the ES8 Ewing sarcoma cell line. While both cell lines represent translocation-driven pediatric mesenchymal tumors, our prior work demonstrated that ES8 cells are resistant to the effects of HSP70 inhibition,^4^ and therefore we chose this as a model to interrogate potential HSP70-independent effects of *DNAJC17*. Cells expressing the dCas9-KRAB chimera were stably transduced with the four different *DNAJC17* deletion alleles of interest. Although expression levels varied, we were able to express all alleles as measured by immunoblot (**Figure 4A**). Further, we confirmed that cellular localization of each allele remained nuclear through live-cell microscopy of GFP-tagged alleles (**Supplementary Figure S3**).

**Figure 4.**
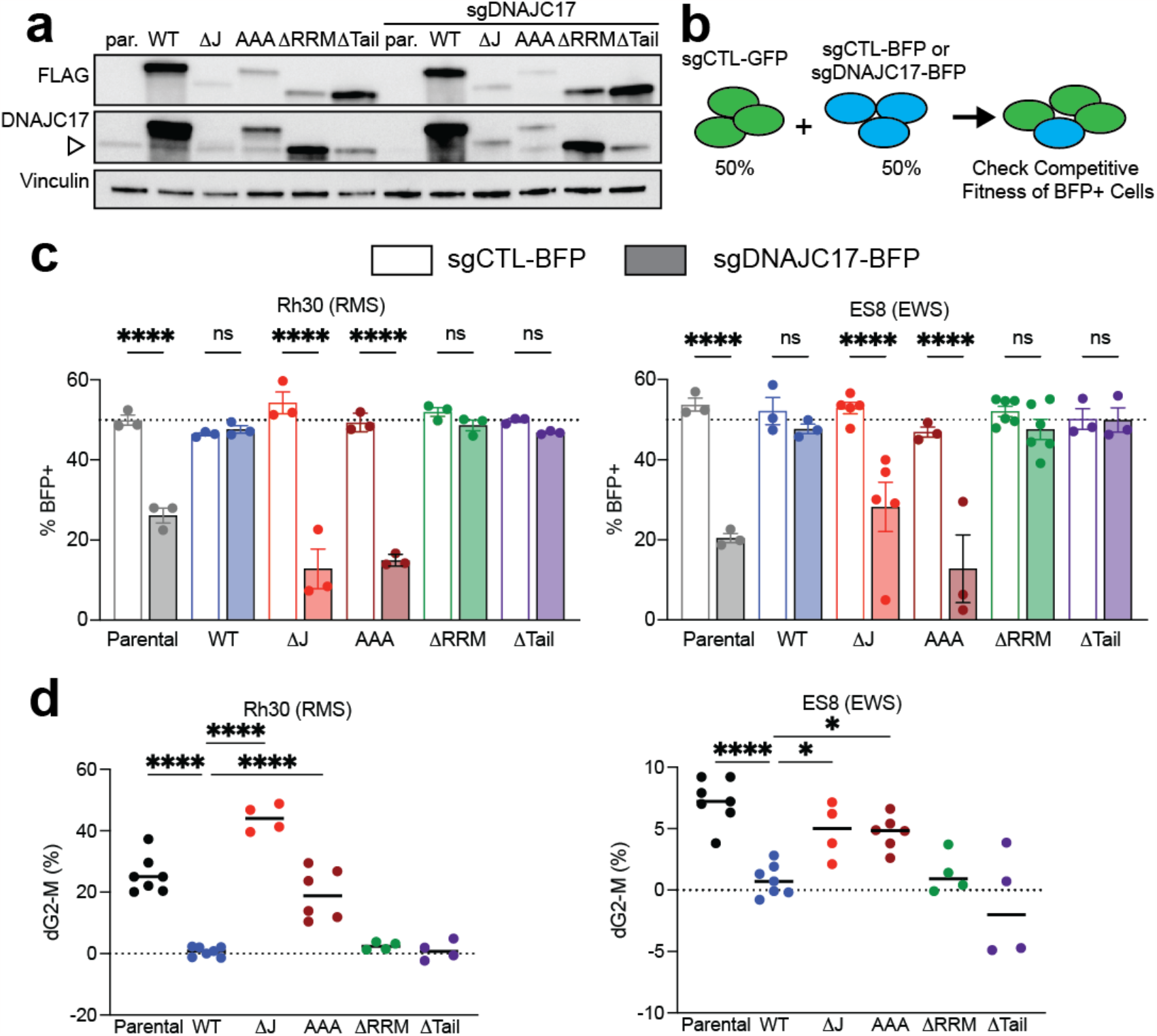
The J-domain but neither the RRM nor the C-terminal tail of DNAJC17 are required for cell cycle progression in cancer cell lines. A, stable expression of the indicated alleles of *DNAJC17* in the Rh30 cell line. Transduction with an sgRNA targeting the 5’UTR of *DNAJC17* results in loss of endogenous (open triangle), but not transduced DNAJC17 protein. B, schematic of competitive fitness assays used to test the ability of *DNAJC17* alleles to rescue cells from endogenous loss. C, results of *DNAJC17* knockdown in cells parental cells, or those expressing the indicated alleles, after four days in culture. Differences significant by 1-way ANOVA; **** p < 0.0001 by post-hoc Holm-Sidak’s test. D, Percentage of cells in G2-M phase following knockdown of endogenous *DNAJC17* shows that wildtype, ΔRRM, and ΔTail can rescue cells from loss of *DNAJC17*. Differences significant by 1-way ANOVA; * p < 0.05 **** p < 0.0001 by post-hoc Holm-Sidak’s test.

Cells were then transduced with lentiviral particles including either non-targeting control sgRNA and a GFP marker, or sgRNA targeting *DNAJC17* immediately 5’ to its transcriptional start site and BFP. This guide RNA design permitted suppression of the endogenous, but not exogenous lentivirally-expressed alleles of *DNAJC17* (**Figure 4A**). Following puromycin selection, equal numbers of cells were plated and the percentage of BFP positive cells was measured over time (**Figure 4B**). In both cellular contexts, we confirmed that while the N-terminal J-domain was essential for cellular survival, the RRM and C-terminal tail of the protein were dispensable **(Figure 4C)**.

To investigate the basis for this attrition of cells, we undertook cell cycle analysis through EdU incorporation assays. Knockdown of DNAJC17 led to a marked reduction in S-phase entry in both Rh30 and ES8 cells, with evidence of G2-M arrest (**Figure 4D**). This could be fully reversed by expression of the ΔRRM and ΔTail, but not the ΔJ mutants. We conclude that DNAJC17 plays a critical role in supporting cellular survival through cell cycle progression, and that a functional J-domain is essential for this.

### DNAJC17 supports normal exon inclusion in splicing events

We next undertook RNASeq of cells following CRISPRi suppression of *DNAJC17* to interrogate the connection between this co-chaperone and cell cycle arrest. Analysis of differentially expressed genes showed a relatively narrow overlap between Rh30 and ES8 cells, with 24 common upregulated and 2 common downregulated genes using a fold change cut-off of 2 (**Figure 5A**). Strikingly, more than three-quarters of these transcriptional changes affected non-protein coding genes (**Table 1**). Interestingly, 5/18 of these genes were small nucleolar RNAs; DNAJC17 has previously been implicated in mRNA splicing based on functional data^16^ and proteomic association with the PRP19 complex.^18,19^ Accordingly, we re-analyzed transcriptome data to identify derangements in splicing by mapping reads across exon boundaries.^20^ Alternative splicing events were readily detectable in the *DNAJC17* knockdown cells, with the majority of these events resulting in exon-skipping in both Rh30 and ES8 cells (**Figure 5B**). RNA-Seq of cells stably expressing the ΔRRM mutant which showed enhanced biochemical activity and full rescue in cellular assays suppressed exon skipping events in both cell lines (**Figure 5C**), suggesting that restoration of normal splicing was associated with rescue of viability. Gene set enrichment analysis^21,22^ of the 24 genes commonly affected by exon skipping events identified enrichment for genes involved in the G2-M checkpoint, further evidence of a molecular connection between splicing and the observed phenotypic consequences of *DNAJC17* knockdown (**Table 2**).

**Table 1.** Transcripts coordinately regulated by *DNAJC17* loss in both ES8 and Rh30 cells.

| RNA Class | Gene | log<sub>2</sub> Fold-Change | RNA Class | Gene | log<sub>2</sub> Fold-Change |
| --- | --- | --- | --- | --- | --- |
| lncRNA | AC004520.1 | 1.40 | snoRNA | SCARNA12 | 1.22 |
|  | AC012615.6 | 1.13 |  | SNORA5A | 1.31 |
|  | AC018630.2 | 1.19 |  | SNORA5C | 1.20 |
|  | AC093525.8 | 2.04 |  | SNORD14E | 3.39 |
|  | AC109322.1 | 1.23 |  | SNORD62A | 2.02 |
|  | AC120057.3 | 1.53 |  | SNORD67 | 1.32 |
|  | AC137932.3 | 1.86 | Protein Coding | BAAT | 1.82 |
|  | AL049844.2 | 1.37 |  | DCAF4L1 | 2.65 |
|  | AL513327.1 | 1.08 |  | OVGP1 | 1.87 |
|  | AP003352.1 | 1.38 |  | PPP1R15A | 1.25 |
|  | CACTIN-AS1 | 1.99 |  | RASL11A | 1.82 |
| Read-through | HSPE1-MOB4 | 3.12 |  | MYO5C | -1.04 |
|  | INO80B-WBP1 | 2.33 |  | DNAJC17 | -2.51 |

**Table 2.**
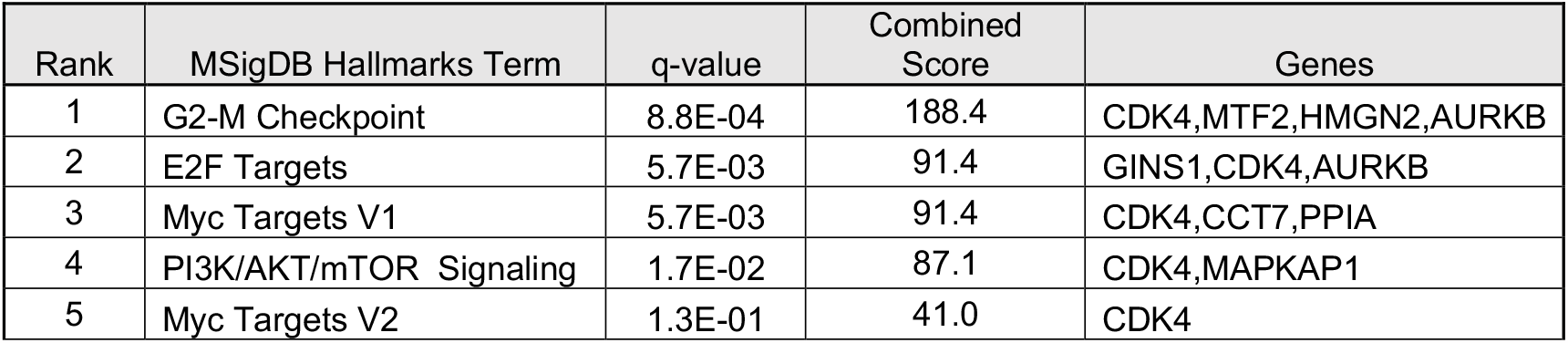
Enrichment analysis of genes with skipped exons upon *DNAJC17* loss in both ES8 and Rh30 cells.

**Figure 5.**
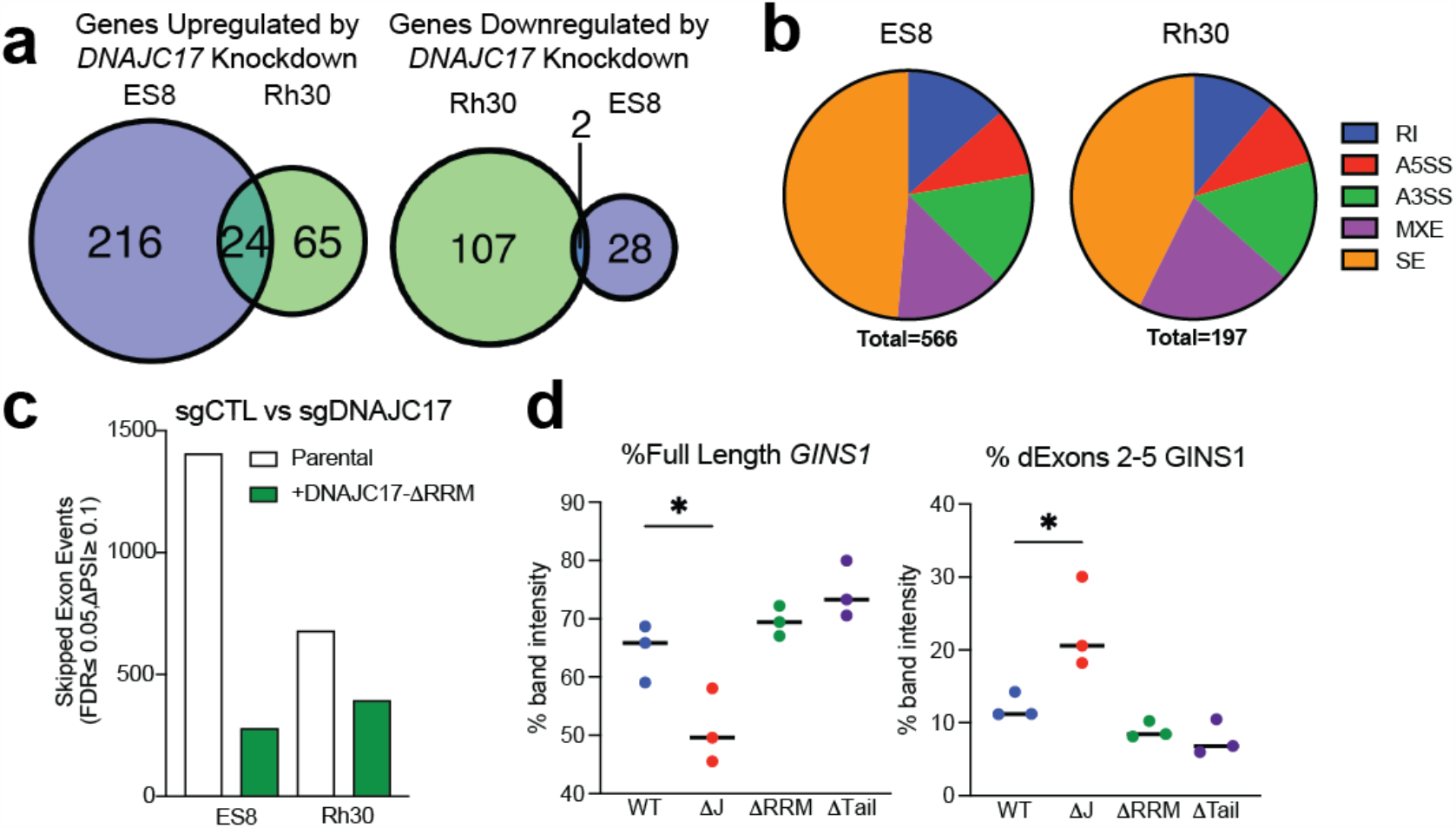
The DNAJC17 J-domain, but not its C-terminal elements, are required for exon inclusion of cell cycle regulators in two human cancer cell lines. A, Overlap of differentially regulated transcripts either up- or downregulated upon *DNAJC17* knockdown in ES8 and Rh30 cells. B, Frequencies of alternate splicing events in ES8 or RH30 cells upon *DNAJC17* knockdown compared to sgCTL cells; RI, retained intron; A5SS, alternative 5’ splicing site; A3SS, alternative 3’ splicing site; MXE, mutually exclusive exon, SE, skipped exon. C, frequency of high-confidence (FDR ≤ 0.05), higher abundance (10% or higher of all transcripts) skipped exons in ES8 or Rh30 cells upon DNAJC17 knockdown. Open bars, parental cells; green bars, cells exogenously expressing DNAJC17-ΔRRM. D, semi-quantitative RT-PCR shows that wildtype, ΔRRM, and ΔTail alleles of *DNAJC17* can rescue correct splicing of *GINS1*, but the ΔJ allele cannot.

*GINS1* encodes the PSF1 component of the GINS DNA replication complex, and transcripts skipping over exon 2 more than doubled upon knockdown of *DNAJC17*. Interestingly, analysis of RMATS data suggested that expression of the ΔRRM allele was completely sufficient to rescue this effect, demonstrating that the RRM is not required for proper regulation of splicing by DNAJC17. We validated and extended this finding from RNASeq to other alleles by semi-quantitative PCR assays. As expected, alleles that rescued RMS and EWS cells from loss of *DNAJC17* showed restoration of full length *GINS1* transcripts. (**Figure 5D, Supplementary Figure S4**).

## Discussion

Evolutionary diversification of the fundamental DnaJ-DnaK machinery found in prokaryotic cells has given rise to 14 different HSP70 and 41 different DNAJ family members in the human genome. Recent work has begun to systematically shed light on the individual functions of different JDP,^18^ which serve not only to provide selectivity to HSP70 but to carry out their own unique activities. Here, we provide additional detail to the cellular functions of a pan-essential JDP through analysis of its component structural domains.

Given the significant structural diversity present in the JDP family, one question that arises is how critical allosteric interactions with HSP70 remain in J-domain protein’s cellular functions. Such requirements may be cell-type or cell-state dependent. For instance, conditions of proteostatic stress create a requirement for the Cwc23 J-domain that is otherwise dispensable, suggesting that this protein may play either a minor or compensatory role in Ssa activation in *S. cereveisae*.^12^ In contrast, we find that the J-domain of DNAJC17 is fundamental to its activity in human cancer cell lines, and that loss of the J-domain yields a protein that cannot support cellular survival. Thus, *DNAJC17* represents a critical adaptation of the HSP70-JDP module in mammalian cells.

Interestingly, purified DNAJC17 does not exhibit HSP70 allosteric interactions, however. We instead find that this function is inhibited by the C-terminal RRM. We hypothesize that the normal regulation of DNAJC17 through RNA binding and/or protein-protein interactions induces an allosteric structural change that relieves this auto-inhibition. Although we were unable to detect RNA-binding by DNAJC17, productive transient interactions between the RRM and RNA may indeed be the basis for this regulation. Additional structural modeling and analysis may help test this hypothesis in the future.

The pan-essential nature of DNAJC17 is at odds with other proteostasis network components such as the 14 different HSP70s, many of which exhibit some degree of redundancy. Others have identified a role for DNAJC17 in regulation of splicing, and we confirm here that knockdown of *DNAJC17* leads to widespread exon skipping events that are enriched in regulators of cell-cycle progression, culminating in G2-M arrest. The molecular connection between DNAJC17 and exon exclusion within this subset of cell cycle regulators is still uncertain, but published proteomic data implicate either the PRP19 complex or components of the U5 spliceosome as being direct interactors of DNAJC17.^16,19^ We did not find evidence that deletion of either the J-domain or RRM alter interaction with components of these complexes (CDC5L or SNRNP200) respectively (data not shown), suggesting that the deletion mutants we generated might alter function but not physical interaction with the relevant splicing machinery.

Together, these data support a model wherein cellular specialization of J-domain function is achieved through localization, protein-protein interactions, and well as regulatory modules (RRM) that may ultimately be dispensable for biochemical function. These structural modifications permit the significant contribution of protein homeostatic machinery to diverse biologic processes such as RNA splicing. In this context, protein chaperone machinery can thus contribute to RNA quality control. Future elucidation of the many roles that HSP70-JDP pairs have evolved to fill will hopefully provide additional details of the contribution of structure to chaperone function.

## Methods

### Cell culture

Cell lines were obtained from the COG Repository (Rh30, ES8), or purchased from ATCC (293T), Genecopoeia (HEK-293) or Takara Bio (Lenti-X). Rh30 and ES8 cells were maintained in RPM-1640, and 293T, HEK-293, and Lenti-X cells were in DMEM. All media was supplemented with 10% FBS and 1x penicillin/streptomycin, and cells were grown in a 37 degree C incubator with 5% CO2. Cells were tested quarterly for mycoplasma and tested to confirm identity by STR analysis twice a year.

### CRISPRi screen

Lentiviral particles were generated from 293T cells transduced with pooled sgRNA libraries as described,^5^ then used to infect Rh30 cells transduced with dCas9-KRAB-BFP and sorted for BFP positivity. Flow cytometry was used to confirm an MOI of approximately 1 forty-eight hours after infection. Cells were selected in puromycin for 72 hours, an aliquot frozen for t_0_ analysis, and the remainder seeded in 500 cm^2^ tissue culture plates at equal density (8×10^6^/plate). Every 3 days, cells were trypsinized, pooled, counted, then re-plated at the same density to maintain 1000x coverage of each sgRNA construct. After 10 population doublings, cells were viably frozen.

Deep sequencing and data analysis were performed as described.^5^ Briefly, genomic DNA was extracted from t_0_ and t_end_ cells using a DNEasy Blood & Tissue kit (Qiagen), digested to enrich for lentiviral integration sites, and sgRNA sequences were amplified by PCR for subsequent sequencing on an Illumina HiSeq. Reads were aligned to the sgRNA library, and fold-change from t_0_ to t_end_ was calculated. A gene-level score was then calculated as the mean of the top three scoring sgRNAs targeting a given transcript.

### Lentiviral transduction

Lenti-X cells were transfected with plasmids of interest, pCMVdR8.91, and pMD2.g using TransIT-LT1 transfection reagent (Mirus) at a 3:1 ratio. Six hours later, ViralBoost reagent (Alstem) was added at 1:500. Seventy-two hours after transfection, viral particles were harvested from the supernatant, filtered through a 0.45 micron PES syringe, and then added to target cells with 6 μg/mL polybrene. In twenty-four hours, selection was started with either puromycin (shRNA or sgRNA constructs) or hygromycin (pLV-EF1-Hygro constructs) for 3 or 10 days, respectively.

### Plasmids and cloning

Wildtype or deletion mutants of DNAJC17 were ordered as gBlocks (IDT) and cloned into pLV-EF1-Hygro using NEBuilder Master Mix (NEB). Codon-optimized sequences of DNAJC17 (WT, ΔJ, ΔRRM, and ΔTail) and DNAJB1 were purchased as gBlocks (IDT) and cloned into pET28b with an N-terminal 6xHis tag followed by a TEV cleavage site. A codon-optimized sequence of HSPA1A was cloned into pET30a with a C-terminal 6xHis tag preceded by a TEV cleavage site. pET28b-TEV-C9R was purchased from AddGene. Small guide RNAs (shown in Supplementary Table S1) were ordered as individual oligonucleotides (IDT), heated at 95 degrees Celsius for 5 minutes in 100 mM K-acetate, 30 mM HEPES-KOH pH 7.4, 2 mM MgCl_2_ and let cool to room temperature to anneal, and cloned into either sgRNA-Puro-T2A-GFP or sgRNA-Puro-T2A-BFP plasmids (a gift of Dr. Jonathan Weissman, Whitehead Institute, MA) after digestion with BlpI and BstXI.

### Transient Transfections

0.6 ×10^6^ HEK-293 cells were plated in 6-well plates and 24 hours later transduced with 0.3 micrograms of 3x-FLAG-tagged DNAJC17 constructs using Mirus transfection reagent at a 3:1 ratio per manufacturer’s instructions. After 3 days, cells were lysed for immunoblot analysis.

### Protein purification

Bacterial expression plasmids were transformed into Lucigen OverExpress C41 cells and induced with 1mM IPTG at 16°C overnight. Bacteria were then pelleted, resuspended in His-binding buffer (50 mM Tris-HCl pH 8, 750 mM NaCl, 10 mM imidazole, 1 mM PMSF) supplemented with Complete protease inhibitor tablets (Roche), and lysed by Dounce homogenization and sonication. Lysates were cleared by centrifugation at 20000 g for 20 min and incubated with pre-equilibrated Ni-NTA resin (Novagen) at 4°C for two hours.

After several washes, protein was eluted from the resin and run on a HiLoad 16/600 Superdex 200 column (Cytiva). Eluted protein was pooled and concentrated using Amicon Ultra Centrifugal Units (MilliporeSigma).

### ATPase assays

ATPase activity was measured by Malachite Green assay (Sigma). Briefly, 12.5 pmol of Hsp72 was incubated with 1 mM ATP in the presence of varying amounts of J-domain protein in Malachite Green Buffer (100mM Tris Base, 20mM KCl, 6mM MgCl2, pH 7.4, 0.01% Triton X-100). After two hours at 37°C, samples were diluted four-fold and mixed with Malachite Green working solution. Color was allowed to develop for 30 minutes at room temperature, and absorbance was then read at 620 nm.

### RNA Immunoprecipitation

RNA immunoprecipitations were carried out using the Imprint RNA Immunoprecipitation kit (Sigma). Briefly, cells were lysed in Mild Lysis Buffer supplemented with protease inhibitor, ribonuclease inhibitor, and DTT on ice for 15 minutes. Lysates were cleared by centrifugation. Lysate was diluted in IP Buffer to a final volume of 500 ul, and 10% was set aside as input.

For FLAG pulldowns, the remaining lysate was incubated with FLAG M2 magnetic beads (Sigma) overnight. For SNRPA1 pulldowns, lysates were incubated overnight with 10 ul of anti-SNRPA1 antibody (Novus), and then with Protein A Magnetic Beads (Sigma) for one hour.

After binding, beads were washed five times with Wash Buffer. Trizol (Fisher) was then added directly to the beads or input aliquot. RNA and protein were sequentially precipitated, washed, and solubilized per the manufacturer’s protocol. RNA was quantified by NanoDrop and loaded abundance measured using a Pico 6000 kit and Eukaryotic Total RNA assay on a BioAnalyzer (Agilent).

### Flow cytometric competitive fitness assays

Following puromycin selection, equal numbers of cells transduced with either GFP or BFP labeled sgRNA constructs were plated in a single well. A sample of this mixed culture was taken at the time of plating, then four days after plating, for flow cytometric analysis. The percentage of BFP+ cells was normalized to the percentage present when mixed cultures were initiated.

### Cell cycle analysis

Cells were analyzed for cell cycle phase using a ClickIT EdU Alexa Fluor 647 kit (Fisher) and FxCycle PI/RNase Staining Solution (Fisher) followed by flow cytometry on an LSR-II (BD) and analysis using FlowJo.

### Fluorescence microscopy

For live-cell imaging, cells were plated in 6-well dishes and transfected with mEGFP-tagged DNAJC17 or deletion mutants using TransIt-LT1 reagent at a 3:1 ratio (Mirus). Cells were then trypsinized and replated on 4-well chambered coverglass (LabTek) coated with poly-D-lysine (Sigma Aldrich). The following day, cells were treated with Hoescht 33342 (Thermo Fisher) for at least 30 minutes and imaged on an OMX-SR microscope with TIRF 60x objective (UCSF Center for Light Microscopy).

### Immunoblots

Cells for immunoblots were lysed in ice cold RIPA buffer with protease and phosphatase inhibitors (Roche). Lysates were quantified using a DC protein assay (Bio-Rad), boiled in 1x Laemmli buffer, and then run on a 4-16% TGX gel. Gels were transferred onto nitrocellulose membrane and blotted from 3-18 hours in primary antibody, washed three times, and then incubated with HRP-conjugated secondary antibody for one hour. Blots were washed thrice, then imaged using ECL reagent (Amersham) on a GelDoc (Bio-Rad). Blots are representative of at least three independent replicates. Quantitation was carried out by ImageJ. The following antibodies were used in immunoblots: FLAG (Sigma #F1804); DNAJC17 (Sigma #HPA040914); vinculin (ProteinTech #66305); HRP-conjugated anti-mouse IgG (Cell Signaling #7076V); HRP-conjugated anti-rabbit IgG (Vector Laboratories #PI-1000-1).

### RNA Sequencing

Cells plated in triplicate were lysed and total RNA collected using an RNEasy kit (Qiagen), with downstream library preparation and sequencing conducted by Genewiz. Libraries were prepared using the NEBNext Ultra RNA Library Prep Kit for Illumina following manufacturer’s instructions (NEB, Ipswich, MA, USA). The sequencing libraries were validated on the Agilent TapeStation (Agilent Technologies, Palo Alto, CA, USA), and quantified by using Qubit 2.0 Fluorometer (Invitrogen, Carlsbad, CA) as well as by quantitative PCR (KAPA Biosystems, Wilmington, MA, USA). Reads were aligned against National Center for Biotechnology Information Build 37 (hg19) of the human genome. Differential expression was conducted using DESeq2.^23^ Analysis of splicing was conducted using rMATS 4.1.2.^20,24,25^

### Semi-quantitative RT-PCR Assays

Total RNA was extracted from cells using an RNEasy kit (Qiagen), and cDNA was synthesized using a SensiFast cDNA kit (Bioline) using manufacturer’s instructions from 250 ng of RNA. The cDNA was diluted 1:4, and 8 uL was added to 400 nm forward and reverse primers (shown in supplementary table S1) and 2x Fast SYBR Green mastermix (Thermo). Thermocycler settings were 95ºC x 10 minutes to denature, followed by 30 cycles of 95ºC x 30 seconds, 60ºC for 902 seconds, and 72ºC for 45 seconds. Reactions were run on a 10% agarose/TBE gel and visualized on a GelDoc (Biorad). Quantitation of band intensities was conducted using ImageJ, in biologic triplicate.

## Acknowledgements

Funding support from the National Institutes of Health (5K08CA218691), the Frank A. Campini Foundation, and the Buster Posey Foundation to A.J.S. Microscopy was performed at the Helen Diller Family Comprehensive Center Laboratory for Cell Analysis, supported by the National Cancer Institute of the National Institutes of Health under Award Number P30CA082103.

## Author contributions

A.J.S., T.G.B., and D.V.A. devised the experiments and interpreted results. A.J.S., D.V.A., K.K., and J.M. conducted experiments and analyzed data. A.J.S. and D.V.A. prepared the manuscript. All authors reviewed, edited, and approved the final manuscript.

## Data availability

All plasmids, modified cell lines, and other experimental reagents are available to investigators upon email request. Data from the CRISPRi screen used to study the growth effects of JDP are available in online supplementary material accompanying our prior publication.^26^ RNA-Seq data from Rh30 and ES8 cells with or without *DNAJC17* knockdown have been deposited in the National Institutes of Health Gene Expression Omnibus, with accession ID GSE235379.

## Additional information

Competing interests: Dr. Bivona is an advisor to Array Biopharma, Revolution Medicines, Novartis, AstraZeneca, Takeda, Springworks, Jazz Pharmaceuticals, Relay Therapeutics, Rain Therapeutics, Engine Biosciences, and receives research funding from Strategia, Verastem, Kinnate, and Revolution Medicines. Dr. Sabnis, Mr. Allegakoen, Ms. Kwong, and Ms. Morales declare no potential conflicts of interest.

