## Supplemental Figures for "The essential chaperone DNAJC17 activates HSP70 to coordinate RNA splicing and G2-M progression"

**Supplemental Information.**

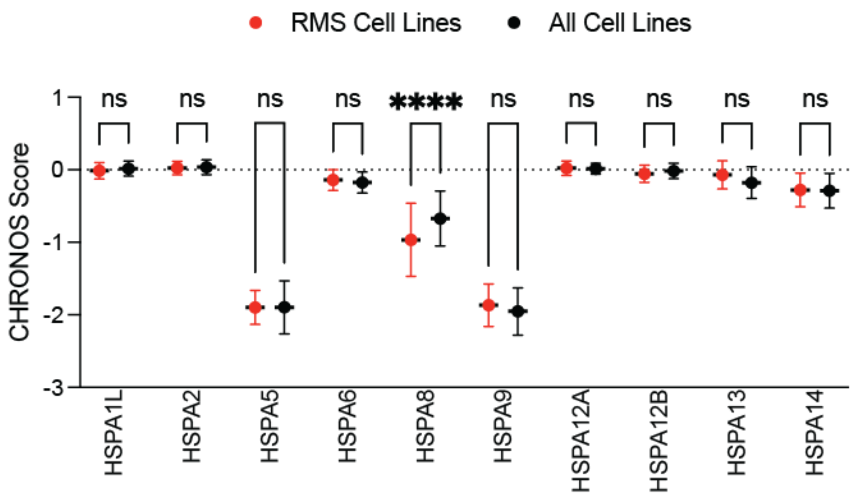


**Figure S1. Cell context-specific proteostasis dependencies uncovered by the Dependency Map.** Analysis of CHRONOS scores shows preferential dependence of rhabdomyosarcoma cell lines (red) on the cytosolic HSP70 encoded by *HSPA8* compared to other cell lines. Mean with standard deviation is plotted. **** p <0.0001 by 2-way ANOVA with post-hoc Holm-Sidak's test.

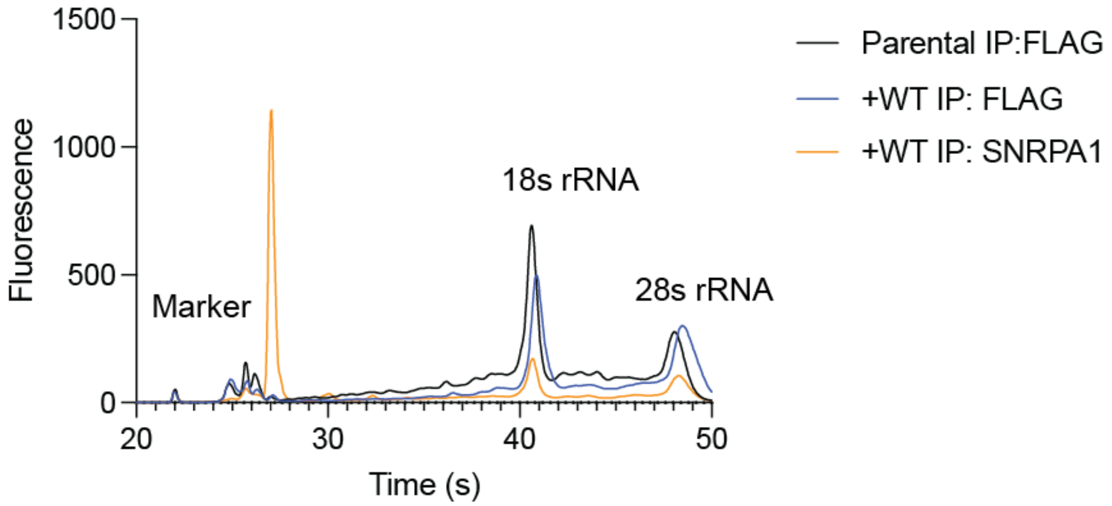


**Figure S2. RNA immunoprecipitation of DNAJC17.** Parental Rh30 cells or cells stably expressing FLAG-tagged *DNAJC17* were lysed, immunoprecipitated with anti-FLAG or anti-SNRPA1 antibodies, and total RNA was extracted. Electrophoretogram shows that while SNRPA1 pulls down small nucleolar RNA as expected, FLAG pulldown does not enrich for any RNAs in FLAG-tagged DNAJC17-expressing cells compared to parentals.

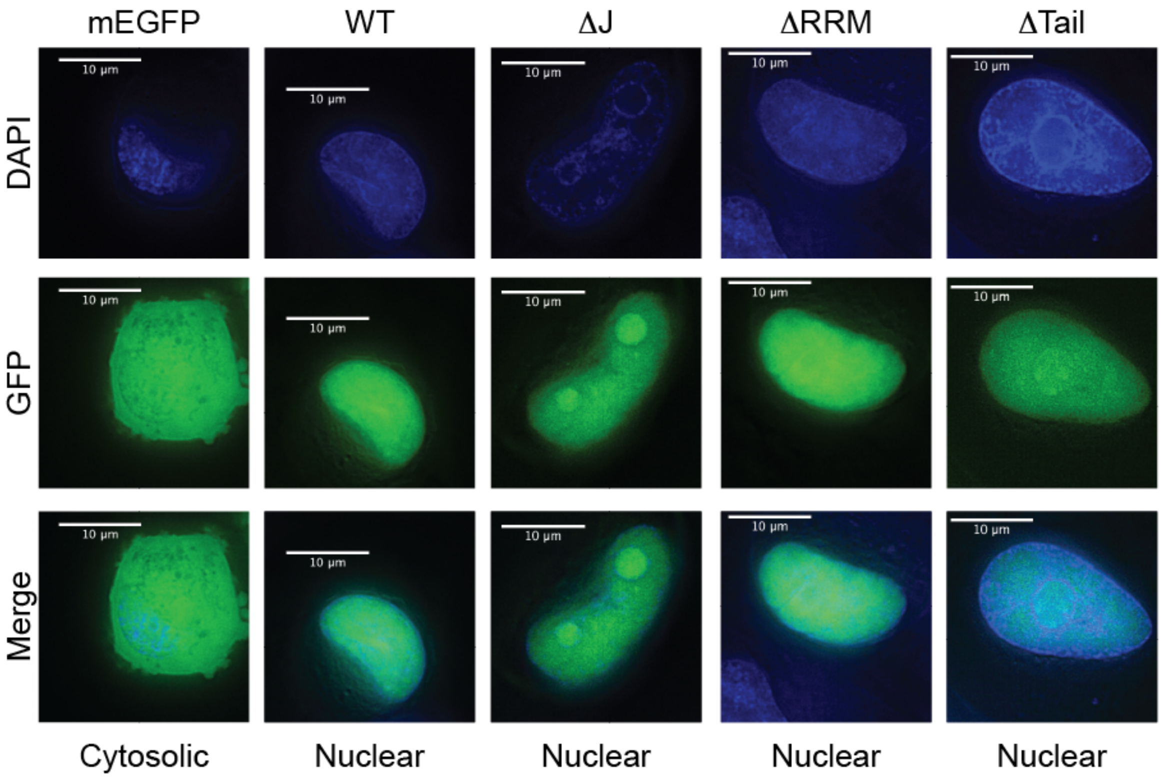


**Figure S3. Deletion of the J-domain, RRM, and C-terminal tail do not alter the sub-cellular localization of DNAJC17.** Rh30 cells were transiently transfected with mEGFP alone or N-terminal GFP-tagged versions of the indicated constructs, then imaged using widefield deconvolution microscopy.

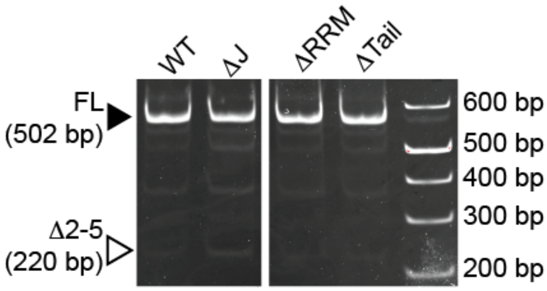


**Figure S4. Semi-quantitative RT-PCR assay to confirm alternative splicing.** Rh30 cells expressing the indicated *DNAJC17* alleles were transduced with sgRNA to knockdown endogenous *DNAJC17*, and total RNA was harvested and used for RT-PCR of GINS1. Gel electrophoresis demonstrates increased abundance of smaller amplicons (open triangle) indicating deletion of exons 2-5 in ΔJ expressing cells, but not others. Quantification of three independent replicates is shown in Figure 6.

**Table S1.** Primers for semi-quantitative reverse transcriptase PCR.

| | |
|---|---|
| *GINS1* 5'UTR Fwd | ATTTTGGCGTGAGAGCTGGT |
| *GINS1* Exon 6 Rev | TCGAGGTAAAAAGTGCTGGCTA |
